# The Integrator complex prevents premature neuronal differentiation through global control of transcription elongation

**DOI:** 10.1101/2024.11.26.625369

**Authors:** Hedda B. Somsen, Kristel N. Eigenhuis, Anne L. Kuijpers-Korporaal, Mariana Pelicano de Almeida, Roberto Montoro Ferrer, Kelsey Kats, Mike R. Dekker, Thuur Zuidweg, Zakia Azmani, Mirjam C.G.N. van den Hout, Wilfred F.J. van IJcken, Danny Huylebroeck, Maarten Fornerod, Raymond A. Poot, Debbie L.C. van den Berg

## Abstract

Transcription pause-release constitutes a key step in the regulation of gene expression. Pathogenic variants in various pausing regulators have been identified in neurodevelopmental disorders, but how transcription pause-release regulates neural development remains largely unknown. Here we show that a patient-specific in-frame deletion in Integrator subunit 8 (INTS8) that prevents association of this subunit with the Integrator complex results in a global increase in nascent transcription and precocious expression of neuronal genes in progenitor cells. Consequently, differentiating neural progenitors fail to sustain a proliferating progenitor cell population during cortical neuronal network formation, resulting in network collapse. We show that attenuation of RNApol2 pause-release by targeted degradation of BRD4 can revert neuronal gene activation and prevent premature progenitor cell loss. Our data demonstrate that a fine-tuned balance between RNApol2 pausing and elongation is crucial for normal neuronal development and suggest transcription pause-release as a druggable target for neurological defects in developmental disorders.

## INTRODUCTION

The cerebral cortex is our information processing centre divided into functional areas that serve higher cognitive functions, including language, memory and thought. It is generated from a diverse pool of progenitor cells that are able to both self-renew and establish the impressive cell diversity of neuronal and glial cell types of the adult brain. Throughout cortical development, an intricate balance between self-renewing and neurogenic divisions ensures enduring presence of neural stem cells while also safeguarding the timely production of neurons and glial cells and diversifying the pool of neural progenitors (reviewed in^1^).

Accurate tissue morphogenesis relies on correctly timed and often synchronous transcriptional responses that can be achieved through controlled release of paused DNA-dependent RNA polymerase II (RNApol2) into productive transcript elongation^2^. This step in the regulation of gene expression involves first the formation of a promoter-proximal paused RNApol2 complex bound by negative elongation factor (NELF) and DRB-sensitivity inducing factor (DSIF), which is then released into the rest of the downstream gene body upon phosphorylation of the RNApol2 C-terminal domain (CTD), NELF, and DSIF. Indeed, transcription elongation has emerged as a key point of control in gene regulation in response to developmental cues^3^. Increased pause duration reduces cell-to-cell variability in gene expression and is an important determinant of transcriptional output^4–6^.

The Integrator complex is an evolutionary conserved 14-subunit complex that regulates RNA processing and gene transcription elongation^7^. Associated with the RNApol2 CTD^8^, Integrator uses its RNA endonuclease activity to cleave nascent transcripts, including uridylate-rich small nuclear RNAs (UsnRNAs)^9^, enhancer RNAs (eRNAs)^10^ and unproductive promoter-proximal transcripts^11^. In addition, Integrator plays a role in recruitment of the super elongation complex (SEC) to stimulate release of paused RNA into productive elongation^12,13^, while its association with protein phosphatase 2A (PP2A) prevents pause-release by removal of SEC-mediated phosphorylation marks from the RNApol2 CTD and the DSIF-subunit SPT5^14–17^.

Biallelic pathogenic variants in genes encoding Integrator subunits INTS1, INTS8, INTS11, and INTS13 cause developmental disorders with overlapping clinical features^18–22^. Intellectual disability (ID), cognitive delay, absent or delayed language development, and typical facial features (e.g., low set ears, bulbous nasal tip, slanted palpebral fissures, abnormal dentition) have frequently been reported, suggesting that Integrator is important for development, including development of the nervous system. The role of Integrator in neural development has been studied in *Drosophila*, where it prevents dedifferentiation of intermediate neural progenitors, and in mice, where acute depletion of Integrator subunits Ints1 and Ints11 results in cortical neuronal migration defects^23,24^. How pathogenic gene variants in Integrator subunits affect neural development in humans remains unclear.

Here, we introduced a homozygous in-frame deletion (ΔEVL) in *INTS8* in human neural progenitor cells (NPCs) to mimic the defective gene product that causes a neurodevelopmental syndrome with cerebellar hypoplasia and spasticity (NEDCHS, OMIM #618572). INTS8^ΔEVL^ does not integrate into the Integrator complex, which compromises its phosphatase activity and causes a global increase in transcription elongation. This aberrant pause-release induces expression of neuronal genes and primes NPCs for differentiation. Premature loss of proliferating progenitors during cortical neuronal network formation provides a rationale for the smaller brain size and associated ID in patients carrying pathogenic variants in INTS8. Treatment of differentiating NPCs with BET Bromodomain degrader dBET6 to inhibit pause-release reverted precocious neuronal gene activation and rescued premature differentiation in INTS8 loss-of-function NPCs. Our findings establish dysregulated transcription elongation as a cause of abnormal brain development and provide a base for further investigating pharmacological targeting of transcription pause-release to ameliorate neurological defects.

## RESULTS

### Pathogenic INTS8^ΔEVL^ variant affects Integrator complex integrity

Bi-allelic gene variants in INTS8 (c.893A>G/c.2917_2925del) cause a neurodevelopmental disorder (NDD) with severe cognitive delay, absent speech, and seizures^20^. Structural brain abnormalities such as microcephaly, cerebellar hypoplasia and nodular heterotopia were also reported. Affected individuals produce INTS8 protein from only one allele, which encodes a variant that harbours a three-amino acid in-frame deletion in its C-terminal region (p.Glu972_Leu974del; referred to as INTS8^ΔEVL^). To investigate the impact of INTS8^ΔEVL^ on neurogenesis, we introduced the variant in human NPCs by CRISPR/Cas9 genome editing (Figure 1A). Two independent homozygous INTS8^ΔEVL^ clones were obtained. The corresponding NPC lines, similar to patients, only produce the INTS8^ΔEVL^ variant protein. We did not detect additional CRISPR edits among the predicted top 10 exonic off-target locations (Figure S1A). Immunocytochemistry (IHC) showed that all lines uniformly express NPC transcription factors SOX2 and PAX6, and the intermediate filament protein NESTIN (Figure 1B). Diffuse staining for the immature neuronal marker beta-III tubulin (TUBB3) with TUJ1 antibody was also observed in our NPC cultures (Figure 1B). Quantitative PCR (qPCR) confirmed expression of neural progenitor marker genes *PAX6*, *SOX2*, *NES* and *MSI1*, as well as proliferation markers *MKI67* and *PCNA* at levels comparable to control NPCs (Figure S1B). INTS8^ΔEVL^ NPCs therefore display stem cell characteristics similar to control NPCs.

**Figure 1.**
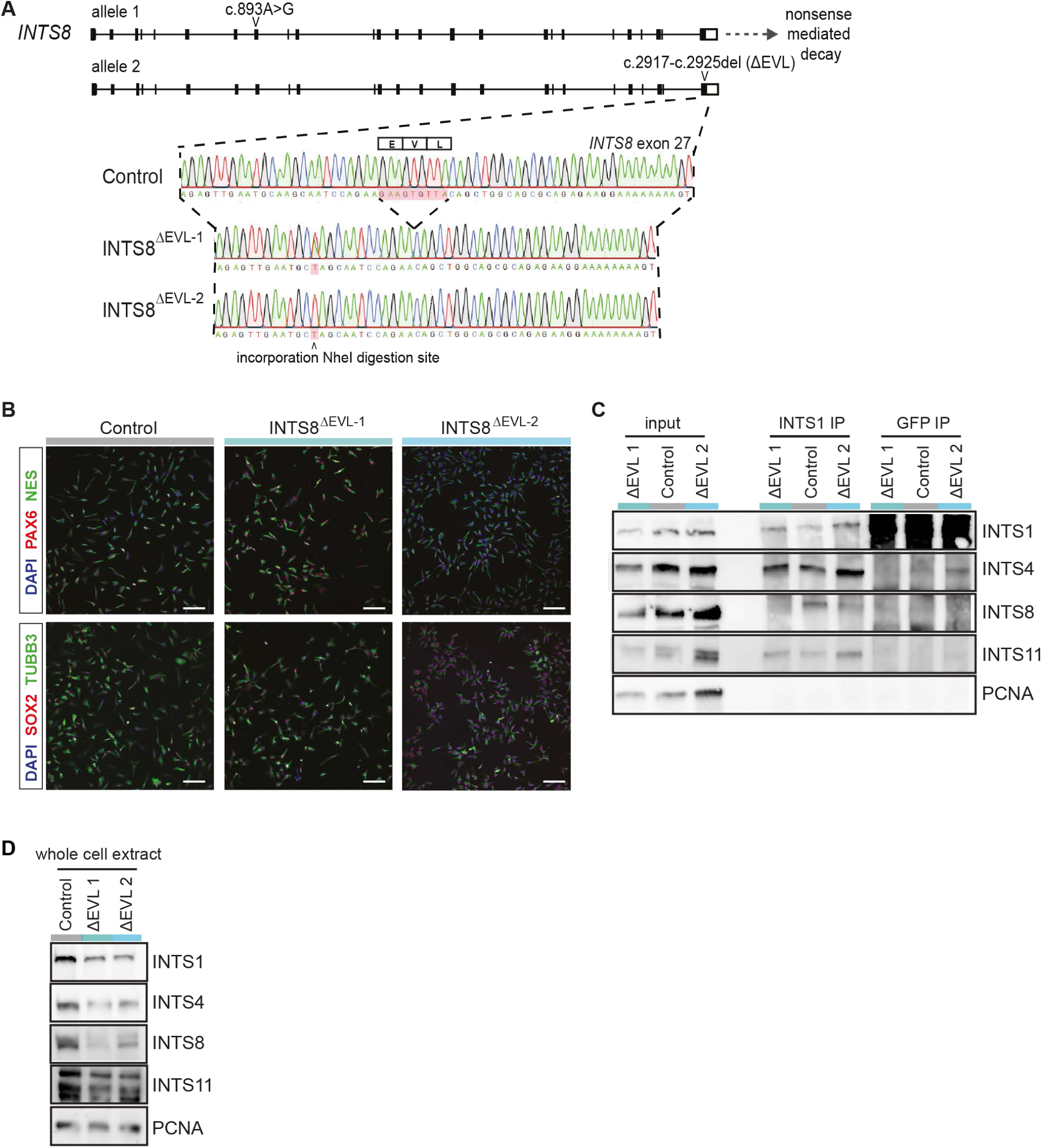
INTS8^ΔEVL^ destabilises the Integrator complex in human NPCs. (A) Generation of NPC lines expressing the pathogenic INTS8 variant. Sanger sequencing trace shows the 9 bp in-frame deletion and incorporated NheI site. (B) Immunocytochemistry characterisation of NPC lines with indicated antibodies. Scale bar represents 50 µm (C) Western blot with indicated antibodies on INTS1 immunoprecipitates from control and INTS8^ΔEVL^ NPC nuclear extracts. GFP antibodies were used as control. (E) Western blot with indicated antibodies on whole cell extracts from control and INTS8^ΔEVL^ NPCs. PCNA serves as loading control.

INTS8, together with INTS5 forms the shoulder module of the Integrator complex and cryo-electron microscopy (cryo-EM) has shown that the EVL amino acid sequence is located at the interface with INTS5, INTS6 and PP2A^17^. Exogenous FLAG-tagged INTS8^ΔEVL^ was previously shown to associate less strongly with selected Integrator subunits, in particular INTS1, INTS12, and the catalytic submodule INTS4/9/11^20,25^. To probe whether Integrator complex incorporation of endogenous INTS8 is indeed affected by the ΔEVL mutation, we immunoprecipitated the complex from control or INTS8^ΔEVL^ NPC nuclear extracts using an antibody against the Integrator core subunit INTS1. Western blot analysis showed that INTS1, INTS4 and INTS11 were purified at comparable efficiency from control and INTS8^ΔEVL^ samples (Figure 1D). In contrast, the amount of INTS8 co-precipitating with INTS1 was severely reduced in extracts from NPCs with only the pathogenic INTS8 variant. This confirms that also endogenous INTS8^ΔEVL^ associates less efficiently with the Integrator complex. Moreover, in line with data from patient fibroblasts^20^, protein levels of INTS8^ΔEVL^ as well as Integrator subunits (i.e., INTS1, INTS4, and INTS11) were reduced in whole-cell extracts (Figure 1E). Importantly, the corresponding transcript levels were not affected (Figure S1C), showing that by failing to integrate into the complex, INTS8^ΔEVL^ likely directly affects protein stability of additional Integrator subunits.

### INTS8-deficient Integrator globally increases nascent transcription in NPCs

Integrator has a widely recognized role in the regulation of transcription of protein coding genes, in particular at the stage of promoter pause-release^12–16^. Mapping of Integrator complex genomic binding sites in our NPCs by ChIP-sequencing using an anti-INTS11 antibody indeed shows that 48.8% of binding sites occur in promoter-proximal regions (Figure 2A). Most of such promoter-proximal binding sites (i.e., 43.6%) are found within 1kb of the transcription start site (TSS), which is also evident from the average peak distribution over scaled genic regions (Figure S2A). The remaining non-promoter associated INTS11 peaks mainly localize to intronic and intergenic regions (Figure 2A).

**Figure 2.**
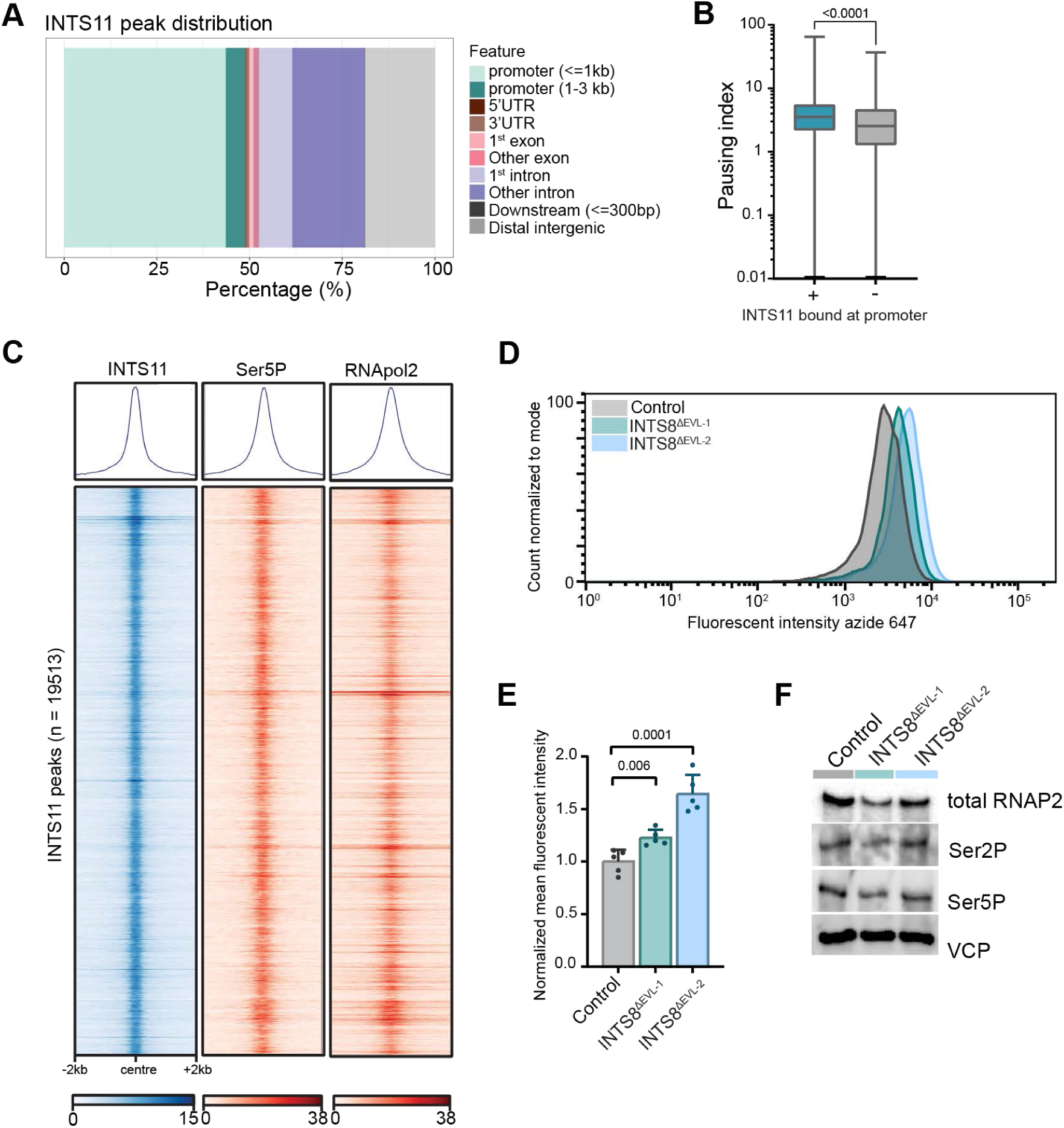
Integrator globally attenuates transcription elongation. (A) Bar graph showing distribution of INTS11 peaks over indicated genomic regions. (B) Boxplot representing distribution of pausing indices for INTS11 promoter-proximal bound genes (n = 6183) versus all other (n=1503) expressed (FPKM>1) genes. (C) Heatmap of 4kb regions centered around 19,513 INTS11 binding sites displaying normalised, sequencing-depth adjusted coverage as determined by ChIP-seq (INTS11) or CUT&RUN (Ser5P and total RNApol2). (D) Representative flow cytometry curves after 1h of 5-Ethynyl-Uridine (EU) incorporation. (E) Normalized mean EU fluorescence intensity from five independent experiments. Error bars represent SEM, p-values determined by t-test, n=5. (F) Western blot with indicated antibodies on whole cell extract from NPC lines.

We then mapped total and paused Ser5-phosphorylated RNApol2 by CUT&RUN. We calculated pausing indices of all expressed (FPKM>1) genes by dividing total RNApol2 length-normalised promoter-proximal reads over gene body reads. This showed that Integrator-bound genes in NPCs on average have a significantly higher pausing index than genes not bound by the Integrator complex in their promoter region (Figure 2B). Indeed, Integrator colocalizes with Ser5-phosphorylated paused RNApol2 in hNPCs (Figure 2C).

The N-terminal region of INTS8 mediates association of the PP2A phosphatase module with the Integrator complex, which counteracts CDK9-mediated RNApol2 pause-release through dephosphorylation of the DSIF-subunit SPT5 and the RNApol2 CTD^15^. Acute depletion of INTS8 was shown to induce pause-release genome-wide^14^. To confirm a role of INTS8 in pause-release in NPCs, we assessed global nascent transcription in the presence of INTS8^ΔEVL^ NPCs by quantifying incorporation of the uridine analog 5-ethynyl uridine (5-EU) into newly synthesized RNA-molecules, using fluorescence flow cytometry analysis. Treatment of NPCs with the CDK9 inhibitor flavopiridol showed that, using this approach and as expected, we mainly detect RNA synthesis by RNApol2 (Figure S2B). During a one-hour pulse with EU, INTS8^ΔEVL^ NPCs incorporated significantly more EU than control NPCs. This could be reverted by addition of flavopiridol (Figure 2D, 2E, and S2C), indicative of a global increase in RNApol2-mediated transcription elongation. Western blot analysis showed that this was not accompanied by a general increase in Ser5 or Ser2 phosphorylation of the RNApol2 CTD (Figure 2F), suggesting that the overall rise in nascent transcription predominantly arises from non-hyperphosphorylated RNApol2. Together, these data show that the three amino acid deletion ΔEVL in INTS8 encodes a loss-of-function variant, resulting in a PP2A-deficient Integrator complex that globally induces transcription pause-release in NPCs.

### INTS8^ΔEVL^ NPCs fail to differentiate into mature neuronal networks in vitro

To probe for the consequences of genome-wide dysregulation of transcription pause-release on neural development, we first analysed the viability and the proliferation in INTS8^ΔEVL^ NPCs. Cell cycle analysis with propidium iodide labelling and univariate modelling showed no differences in cell cycle phase contribution between control and INTS8^ΔEVL^ NPCs (Figure S3A). Immunolabelling of NPCs with anti-cleaved caspase 3 and anti-p21 (CDKN1A) antibodies, as indication of apoptosis and cell cycle arrest/senescence, respectively, also did not show significant occurrence of these events in any of the conditions tested (Figure S3B).

Next, we evaluated the ability of control and INTS8^ΔEVL^ NPCs to form neuronal networks in vitro. We adopted a cell culture method that supports the formation of electrophysiologically mature neuronal networks from pluripotent stem cell-derived NPCs in 8-10 weeks (Figure 3A)^26^. In the first week of differentiation, control NPCs develop multiple projections that stain positive (+) for TUBB3 (Figure 3B). In the second and third week of differentiation, MAP2^+^ dendritic and SMI312^+^ axonal projections could be observed in neuronal networks from control cells, indicative of progressive neuronal maturation accompanied by a gradual reduction in dividing KI67^+^ progenitor cells and the acquisition of cortical neuron identity (FOXG1, Figures 3B and S3C). TUBB3^+^ projections similarly develop in differentiating INTS8^ΔEVL^ NPCs. However, we noted a rapid decrease in INTS8^ΔEVL^ cell density under differentiating conditions, resulting in a complete loss of viable cells between D8-D12 of neuronal differentiation. Moreover, compared to control NPCs, morphological changes such as narrowing of the cell body, development of thin processes and organisation into rosette-like structures, appear to be accelerated INTS8^ΔEVL^ variant cells (Figure 3B).

**Figure 3.**
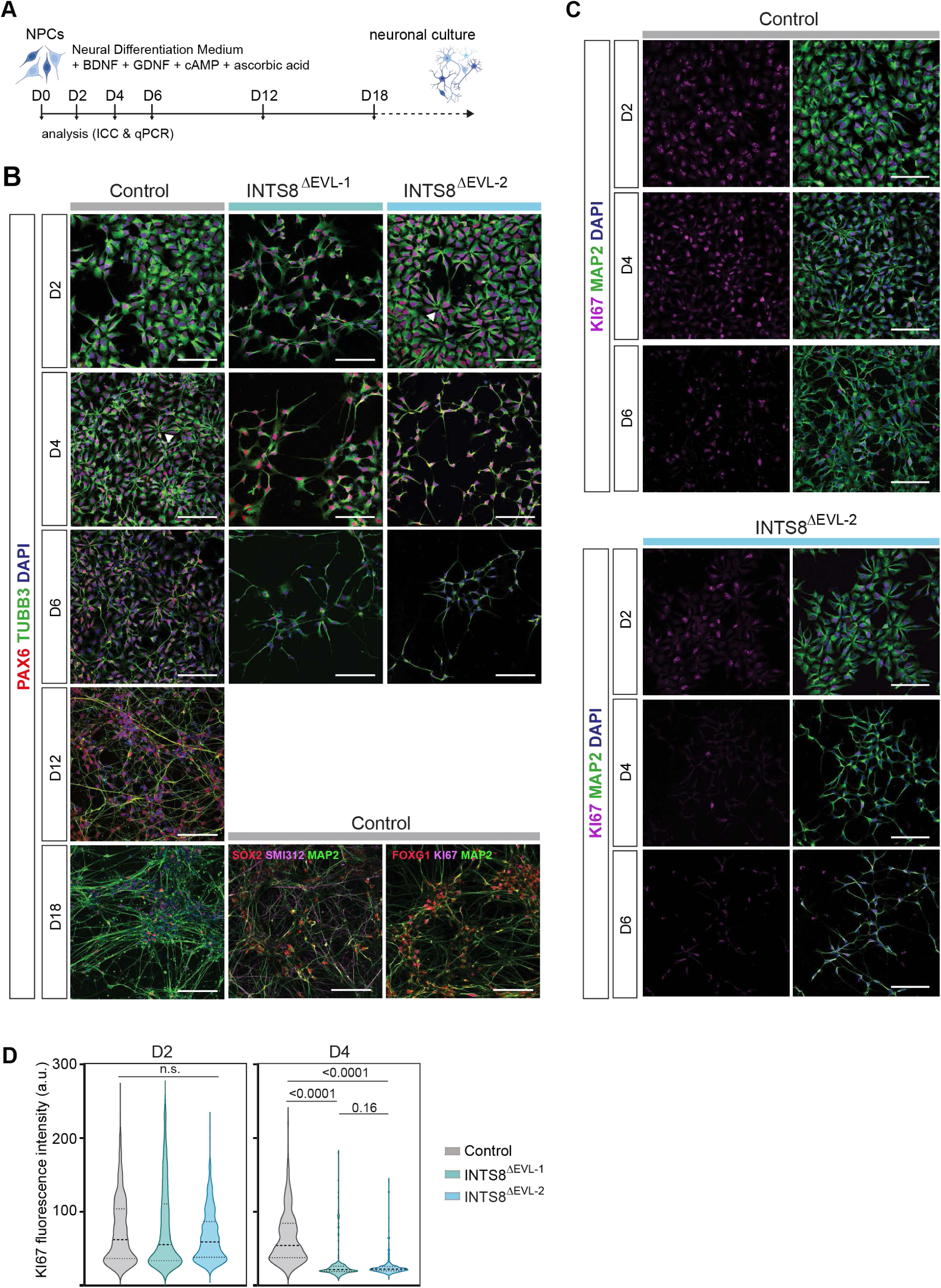
Premature progenitor cell loss and defective neuronal differentiation in INTS8^ΔEVL^ NPCs. (A) Schematic overview of neural differentiation protocol and experimental endpoints. (B) Differentiating NPCs subjected to immunocytochemistry with indicated antibodies analysed at different experimental endpoints. Scale bar 100 µm. (C) Differentiating NPCs subjected to ICC with KI67 and MAP2 antibodies at indicated timepoints during neuronal differentiation. Scale bar represents 100 µm. (D) Quantification of KI67 fluorescence intensity at D2 and D4 of in vitro differentiation. P-values from Mann-Whitney test.

Throughout neural differentiation, self-renewing KI67^+^ progenitors remain present in the population to start generating different neuronal and then glial cell types (Figures 3C and S3C). However, upon withdrawal of mitogens and initiation of differentiation, we observed a rapid decline in proliferating KI67^+^ cells in both of our INTS8^ΔEVL^ lines that was not observed in differentiating control NPCs (Figures 3C and S3D). Quantification of *MKI67* expression and KI67 fluorescence intensity during the first four days of differentiation confirmed this observation (Figures 3D and 5C). These results show genome-wide dysregulation of transcription pause-release together with a compromised proliferative stem cell population during neural differentiation, which ultimately affects neuronal survival and adherent neuronal network formation.

### Precocious neuronal gene expression drives premature differentiation

To identify the pathways that contribute to the observed neuronal differentiation defect in INTS8^ΔEVL^ NPCs, we carried out a genome-wide transcriptome analysis and analyzed the effect of the INTS8^ΔEVL^ variant on gene expression and RNA splicing. Altered splicing patterns, in particular exon skipping, linked to UsnRNA 3’ end processing defects were reported in INTS8^ΔEVL^-variant expressing patient fibroblasts^20^. We did not detect UsnRNA 3’ end processing defects (Figure S4A) and with DEX-seq^27,28^ identified 15 differentially expressed exons (|Log2FC ≥ 2| and p_adj_ ≤ 0.01), belonging to 12 unique genes (Figure S4B). These data suggest that in NPCs, INTS8^ΔEVL^ does not overtly affect the 3’ end processing endonuclease type function of the Integrator complex.

Next, we documented the effect of the INTS8^ΔEVL^ variant on gene expression. In total 328 deregulated genes were identified (|Log2FC|>0.6 and FDR<0.05), 154 of which were up- and 174 were downregulated in the presence of the ΔEVL variant (Figure 4A). For 56% (n=184) of these differentially expressed genes (DEGs), significant Integrator complex binding was observed in the promoter-proximal region (Figure 4B). We performed gene set enrichment analysis (GSEA)^29^ to associate human phenotypes to our NPC transcriptome data. Notably, this GSEA enriched for terms that reflect clinical characteristics of individuals with biallelic variants in *INTS8* (Figure 4C and 4D). These features included ID, but also more specific terms related to human phenotypes (i.e., typical absence or non-motor seizure, gray matter heterotopia) that are present in INTS8 but not INTS1 variant-carrying individuals^18,20^, suggesting that the transcriptional profile of INTS8^ΔEVL^ NPCs captures clinically relevant pathways.

**Figure 4.**
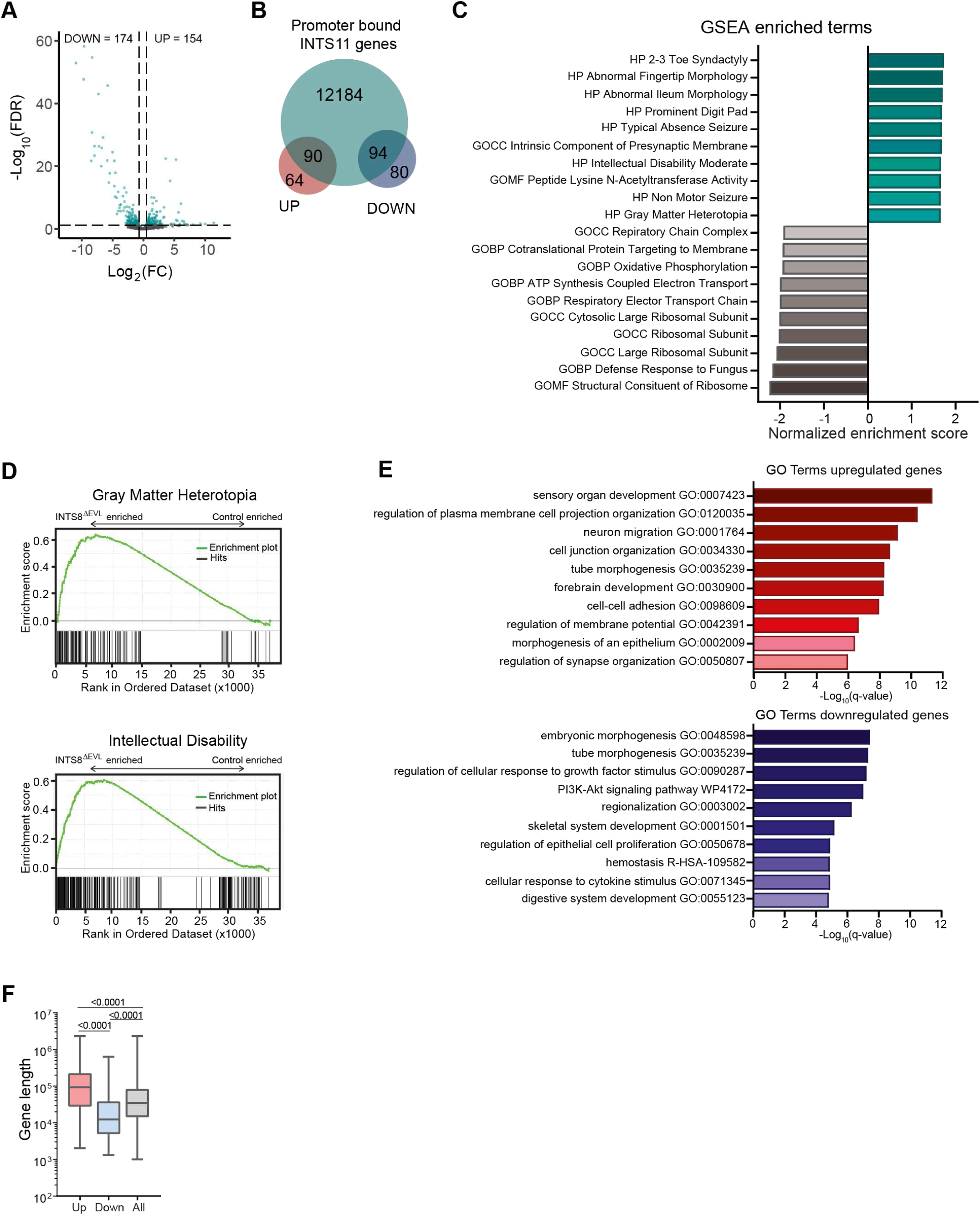
Transcriptomic changes in INTS8^ΔEVL^ NPCs reflect clinical characteristics. (A) Volcano plot visualizing DEGs (green dots) as determined by RNA-seq (|Log_2_FC>0.6|, FDR <0.05).(B) Venn diagram showing overlap between INTS8^ΔEVL^ DEGs and genes with INTS11 bound at the promotor site (TSS+/-500bp) as determined by INTS11 ChIP-seq. (C) Top terms from gene set enrichment analysis (GSEA) terms on control (grey) and INTS8^ΔEVL^ (green) enriched genes. (D) Enrichment score plots from GSEA human phenotype terms Gray Matter Heterotopia and Intellectual Disability. (E) Metascape GO analysis on genes upregulated (red) and downregulated (blue) in INTS8^ΔEVL^ compared to control NPCs. Enriched terms are ranked by Benjamini-Hochberg adjusted p-value (q-value). (F) Boxplot representing gene length (bp) distribution of DEGs versus general distribution. P-values determined by Mann-Whitney test.

**Figure 5.**
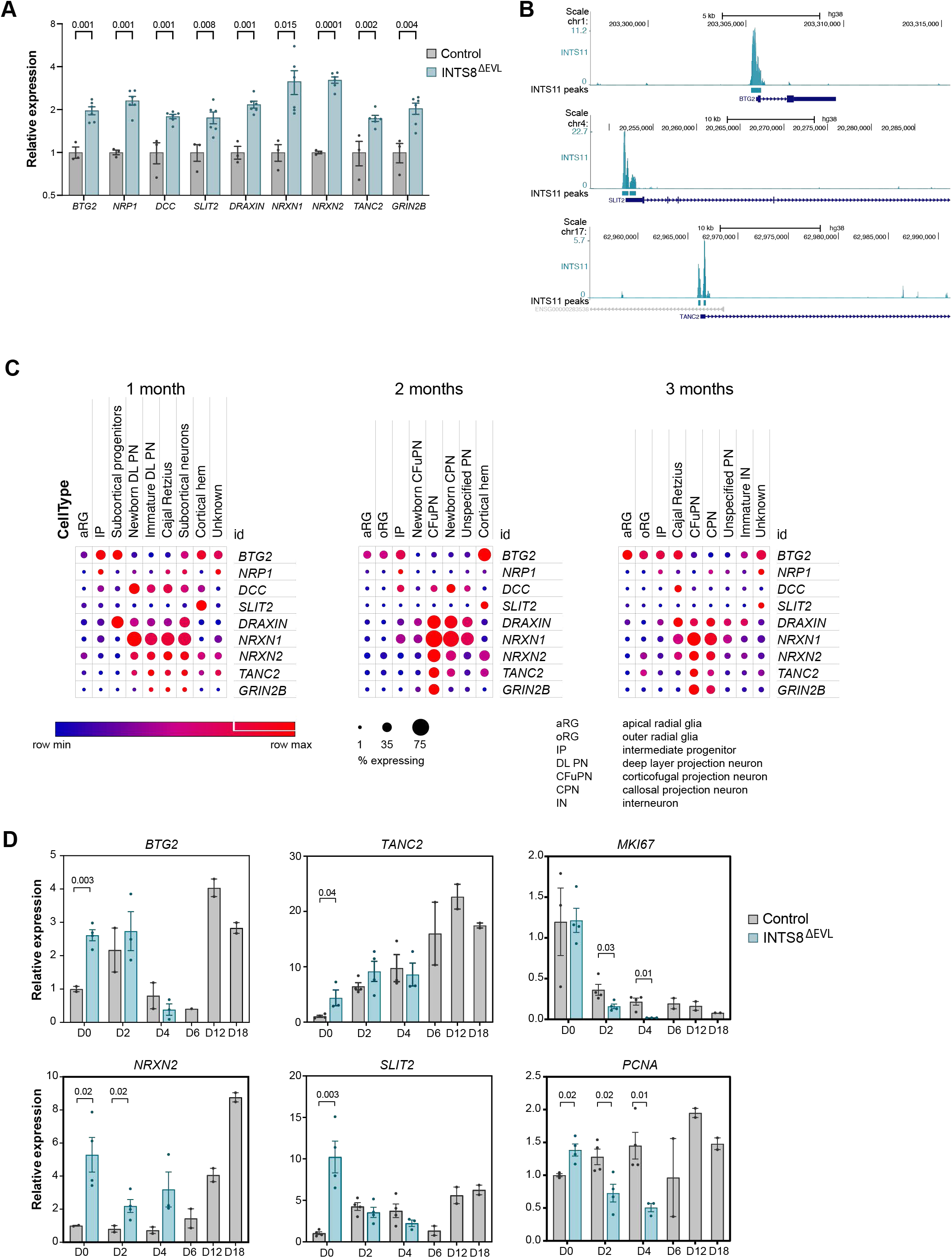
Integrator directly suppresses neuronal gene expression in NPCs. (A) Relative mean expression of neuronal genes in control and INTS8^ΔEVL^ NPCs. INTS11 promoter binding is indicated. Error bars represent SEM, q-values determined by two-stage step-up method of Benjamini, Krieger and Yekutieli, n=3 for control, n=6 for INTS8^ΔEVL^. (B) UCSC browser tracks displaying normalized INTS11 ChIP-seq read density at indicated genomic regions. Significant called peaks are indicated on a separate line. (C) Dot plot representing expression levels of indicated genes in each cell cluster at different stages of cortical brain organoid maturation. Dot colour represents scaled mean expression level across the cluster, while the size of each dot corresponds to the percentage of cells in the cluster expressing the gene (percent expressing). (D) RT-qPCR analysis showing relative mean expression level of indicated genes at different stages of neural differentiation. Error bars represent SEM, q-values determined by two-stage step-up method of Benjamini, Krieger and Yekutieli, n=4 for D0-D4, n=2 for D6-D18.

We then performed gene ontology (GO) analysis on the set of DEGs to identify biological pathways that could explain the progenitor cell loss and accelerated morphological changes in our in vitro neural differentiation system. Indeed, in downregulated genes, we obtained enrichment for terms related to cell proliferation (e.g., regulation of cellular response to growth factor stimulus, PI3K-AKT signalling pathway, and regulation of epithelial cell proliferation). In upregulated genes in INTS8^ΔEVL^ NPCs, GO terms related to forebrain development, cell morphology, cell adhesion, neuronal migration and synapse organisation were enriched (Figure 4E). Similar terms were enriched when only Integrator promoter-proximal bound DEGs were considered (Figure S4C). Furthermore, we noted that the average gene length of upregulated genes was significantly longer than the total average gene length (Figure 5H). This indicates that in INTS8^ΔEVL^ NPCs, released RNApol2 elongation complexes can be sufficiently processive to support the production of long transcripts. Neuronal genes are overrepresented among these long transcripts and potentially contribute to the premature differentiation of INTS8^ΔEVL^ NPCs^30^.

To explore the impact of INTS8^ΔEVL^-induced pause-release on neuronal gene expression and differentiation in more detail, we focused on a subset of upregulated genes expressed at low-to-intermediate levels (0.13<FPKM<23) in control NPCs that contain Integrator binding sites in their promoter region (Figures 5A, 5B, and S4D). First, we analyzed the expression of *BTG2* (*PC3*, *TIS21*), which is largely confined to progenitors in the developing mouse brain and in human cortical organoids (Figure 5C)^31,32^. *BTG2* is transiently expressed as these progenitors commit to neuronal differentiation^33,34^. We performed qPCR analysis at different stages of neuronal differentiation and could detect upregulation of BTG2 in cycling INTS8^ΔEVL^ NPCs, emulated by a temporary increase in expression in control cells at the onset of neuronal differentiation (Figure 5D). Upregulated *Btg2* expression in neocortical progenitor zones of mouse embryos results in microcephaly due to premature onset of symmetric, consumptive NPC divisions^35^. Its untimely expression in INTS8^ΔEVL^ NPCs therefore offers a possible explanation for the observed progenitor cell loss during in vitro neuronal differentiation.

Following up on the anti-proliferative function of BTG2, we examined a set of neuronal genes whose expression was consistently increased in INTS8^ΔEVL^ NPCs (Figure 5A). These included short- and long-range guidance molecules of the Semaphorin (i.e., *NRP1*), Netrin (i.e., *DCC*, *DRAXIN*) and Slit-Robo (i.e., *SLIT2*) signalling pathways involved in neuronal migration and axonal pathfinding, as well as factors involved in synaptogenesis (i.e., neurexins *NRXN1* and *NXRN2*, pre- and postsynaptic scaffolding subunit *TANC2*, and the NMDA receptor subunit encoding gene *GRIN2B*). Analysis of single cell RNA-seq data from human cortical organoids showed that expression of most of these genes is strongly enriched in postmitotic neuronal cell populations at different stages of organoid differentiation (Figure 5C). Assessment of *NRXN2, TANC2* and *SLIT2* transcript levels during in vitro neural differentiation indeed showed that these are upregulated in INTS8^ΔEVL^ NPCs and induced in control NPCs at later timepoints during differentiation (Figure 5D). Furthermore, in line with our IHC characterisation, we observed a significant reduction in transcript levels of proliferation markers MKI67 and PCNA in INTS8^ΔEVL^ NPCs at D2 and 4 of in vitro differentiation (Figure 5D). Taken together, our data suggest that in the presence of INTS8^ΔEVL^, PP2A-deficient Integrator results in aberrant pause-release that induces untimely expression of differentiation-inducing and neuronal maturation genes, which may prime NPCs for neuronal differentiation.

### Inhibition of pause-release rescues INTS8^ΔEVL^ gene signature and progenitor cell loss

So far, our observations of aberrant pause-release and premature neuronal differentiation of INTS8^ΔEVL^ NPCs are correlative. To probe for a causal relation between these events, we tested if the premature progenitor cell loss linked to aberrant pause-release in INTS8^ΔEVL^ NPCs could be restored by inhibiting the activity of CDK9. Indeed, PP2A-Integrator and CDK9 target the same residues in the RNApol2 CTD and SPT5 to regulate the balance between transcription pausing and elongation^15–17^. Inhibiting the CDK9 activity may therefore restore the transcription pausing/elongation balance in INTS8^ΔEVL^ NPCs. To test this hypothesis, we used the BET degrader dBET6^36^, which induces BRD4 degradation and causes a genome-wide induction of transcriptional pausing by inhibiting BRD4-CDK9 activity^36–38^.

We first treated NPCs with different concentrations of dBET6 and monitored the effect on BRD4 protein levels. We observed a dose-dependent reduction in BRD4 protein between 2.5-25 nM dBET6 and showed that a partial reduction in BRD4 levels could be stably maintained for up to 48 hours (Figures S5A and S5B). Moreover, partial BRD4 depletion did not cause a global decrease in the hyperphosphorylated IIo-fraction of the largest RNApol2 subunit RPB1, suggesting there is no global shutdown of transcription elongation (Figures 6A and S5C). In line with these observations, monitoring nascent transcription with 5-EU demonstrated that, compared to the complete transcription elongation block by a 1-hour treatment with 1 µM flavopiridol, low-to-intermediate doses of dBET6 had a moderate effect on global transcription (Figure S5D). Collectively, these experiments show that careful dosing of dBET6 can be used to modulate transcription elongation.

**Figure 6.**
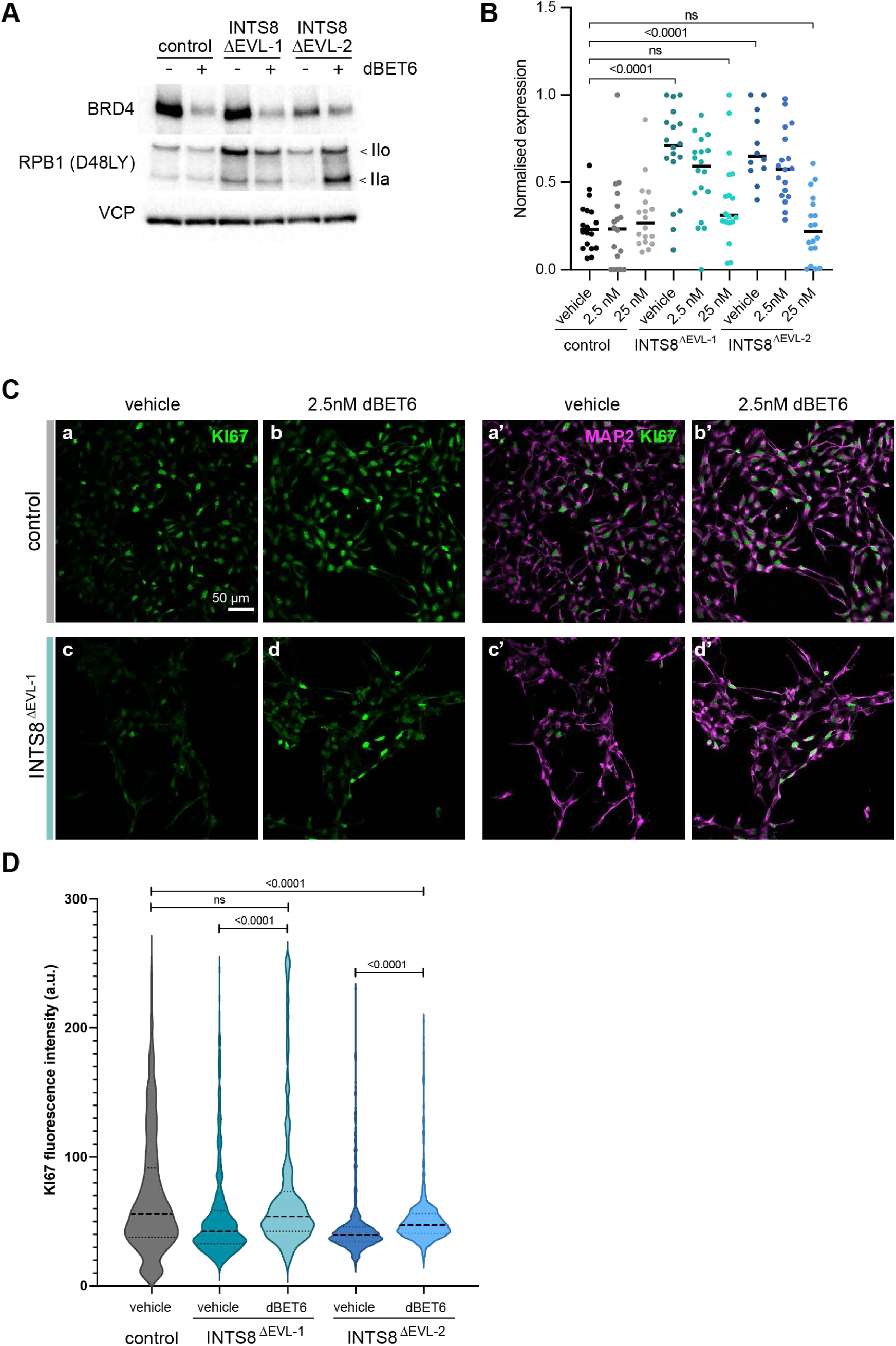
Inhibition of pause-release restores gene expression and rescues premature progenitor cell loss. (A) Western blot with indicated antibodies on whole cell extracts from NPCs treated with vehicle or 25 nM dBET6 for 24 hrs. IIo – hyperphosphorylated, IIa – hypophosphorylated RPB1. VCP serves as loading control. (B) Z-score normalised mean gene expression for *BTG2*, *GRIN2B*, *DRAXIN*, *DCC*, *NRP1*, *SLIT2*, and *NRXN2* in NPCs following 24 hr treatment with vehicle or increasing concentration dBET6. P-values determined by Mann-Whitney test. (C) Immunocytochemistry with indicated antibodies on NPCs at neural differentiation D4, treated with vehicle or 2.5 nM dBET6 for 48-96 hours. Scale bar represents 50 µm. (D) Violin plot representing nuclear KI67 fluorescence intensity at D4 of neural differentiation in cells treated with vehicle or 2.5 nM dBET6 for 48-96 hours.

Next, we tested whether inhibition of CDK9 activity by low-dose dBET6 treatment would be able to revert the increased neuronal gene expression in INTS8^ΔEVL^ NPCs. We treated cells for 24 hours with dBET6 and performed spike-in qPCR analysis for *BTG2*, *GRIN2B*, *DRAXIN*, *DCC*, *NRP1*, *SLIT2*, and *NRXN2*. While the tested concentrations dBET6 did not affect gene expression after 24 hours in control NPCs, we detected a dose-dependent reduction in neuronal gene expression in INTS8^ΔEVL^-producing NPCs (Figure 6B). This confirms indeed that increased pause-release is responsible for upregulated neuronal gene expression in INTS8^ΔEVL^ NPCs. We then subjected differentiating NPCs to dBET-treatment for 48-96 hours and monitored progenitor cell presence in the population by IHC, using anti-KI67 antibody. While treatment with 10 and 25 nM dBET6 during this period prevented neuronal differentiation of both control and INTS8^ΔEVL^ NPCs (Figure S5E), normal differentiation occurred in the presence of 2.5 nM dBET6, as shown by IHC with an antibody against dendritic marker MAP2 (Figure S5E and S5F). Compared to vehicle-treated controls, we observed a significant increase in mean KI67-fluorescence intensity across both INTS8^ΔEVL^ populations in two biological replicates treated with 2.5 nM dBET6 (Figures 6C, 6D, and S5F). We conclude that a correctly tuned balance between PP2A-Integrator-stimulated transcriptional pausing and CDK9-stimulated pause-release is crucial for accurate timing of cell-fate decisions during neuronal differentiation. Moreover, our data provide evidence for the pharmacological targetability of this balance to potentially ameliorate neurological defects.

## DISCUSSION

By using a previously identified pathogenic variant in Integrator subunit INTS8, and its functional complementation by the inhibitor dBET6, we show for the first time that the balance between transcription pausing and elongation correlates with precise orchestration of cell fate decisions during human neurodevelopment. Transcription pause-release is a fundamental process that contributes to greater precision in timing and levels of gene expression^2,3,5^ and enables rapid, synchronous responses to external cues^39–41^. We show that in NPCs, the Integrator complex binds the promoter regions of neuronal genes and attenuates their expression by counteracting pause-release through the INTS8- associated phosphatase module. This suggests that dampening neuronal gene expression by transcriptional pausing is crucial to maintain a progenitor pool during neural differentiation. Failure to do so provides a rationale for the microcephaly phenotype observed in individuals that exclusively produce the INTS8^ΔEVL^ variant^20^.

Besides Integrator subunits, pathogenic variants in various additional genes encoding regulators of transcription pause-release (e.g., Super Elongation Complex subunits^42–46^, PAF1 complex subunits^47^, and BRD4^48,49^) have been reported to cause NDD, underscoring the importance of accurate transcription elongation control for normal brain development and its clinical relevance. Importantly, we demonstrate that inhibition of CDK9-activity can compensate for the functional compromise of Integrator’s phosphatase activity, restore precocious neuronal gene expression, and prevent progenitor cell loss during neural differentiation. Antagonistic activities of CDK9 and PP2A-Integrator in regulating the balance between RNApol2 pausing and elongation have been shown in cancer cells, where CDK9 inhibitors act synergistically with small molecule activators of PP2A to enhance transcription pausing and thereby tumor cell killing^16^. We show that modulation of this balance could also be considered as a therapeutical strategy in certain types of NDD. Specifically, during postnatal brain development and maturation, paused RNApol2 release drives the induction of immediate early gene (IEG) expression in response to neuronal activity and as such plays an important role in neuronal plasticity and learning^50,51^, extending the potential therapeutic window beyond the foetal phase. Pharmacological modulation of pause-release in NDD potentially benefits from the various compounds that are being developed in (pre-)clinical studies to target transcription elongation in cancer^16,52^ or HIV-1^53,54^ infected patients.

Acute INTS8 depletion by RNA interference or degron-based protein degradation results in a global release of RNApol2 into gene bodies and upregulation of >1000 genes^14,15^. We identified a total of 328 DEGs with no clear bias towards up- or downregulated gene expression and did not detect RNApol2-CTD hyperphosphorylation. Compensatory mechanisms likely come into play during long-term INTS8 loss-of-function, but whether the PP2A holoenzyme can be targeted to paused RNApol2 independently of Integrator remains unclear. Alternative candidate phosphatases that could take over the role of PP2A-Integrator in catalysing dephosphorylation of SPT5 and the RNApol2-CTD to dampen pause-release are PP1 and PP4. PP1 was reported to immunoprecipitate with INTS6 and synergize with PP2A in mediating dephosphorylation of substrates of CDK9^16^. PP4 has been shown to bind to promoter-proximal regions and target phosphorylation sites of CDK9 in SPT5, including the Ser666 residue also targeted by PP2A-Integrator^55^. However, given the profound developmental phenotype of the *INTS8^ΔEVL^* gene variant, PP1 and PP4 unlikely are fully complementary.

In addition to the INTS8 variant investigated in this study, genetic variants in INTS1, INTS11 and INTS13 have also been described to cause NDD^18–21,56^. While the clinical features of these disorders largely overlap, the affected subunits belong to different Integrator submodules and play distinct roles in the regulation of transcription elongation. Where INTS1 and INTS8 form part of the eight-subunit core of Integrator, the endonuclease INTS11 constitutes the catalytic subunit of the cleavage module and is contacted by the tail module subunit INTS13 to promote termination of promoter-proximal paused RNApol2^57,58^. Similar to INTS8^ΔEVL^, INTS11 and INTS13 proteins that harbour pathogenic missense variants associate less strongly with the Integrator core complex and RNApol2, indicating that a loss-of-function mechanism is at play here as well^19,22^. Destabilisation of the Integrator complex, including of its INTS4/INTS9/INTS11 endonuclease module, in INTS8^ΔEVL^-expressing NPCs as well as in patient fibroblasts^20^ provides a rationale for the reported similarities in clinical presentation. Further identification of common developmental pathways and neurological functions affected by dysregulated transcription elongation, as well as the broader applicability of pharmacologically targeting of this process, may be an important area of future study.

## MATERIALS AND METHODS

### Cell culture

GIBCO H9 hESC-derived neural stem cells (N7800) were cultured at 37⁰C with 5% CO_2_ on plates coated with 75µg/ml Geltrex™ LDVE-Free Reduced Growth Factor Basement Membrane Matrix (Gibco A1413292). Cells were maintained in KnockOut™ DMEM/F-12 (12660012), 2% StemPro Neural Supplement (A1050801), 20ng bFGF/ml (Peprotech 100-18B), 20ng EGF/ml (Peprotech 315-09) and 2mM L-Glutamine (Gibco 25030-024). Cells were passaged with accutase (Sigma A6964).

### Antibodies

For immunoprecipitation and Western blotting, primary antibodies raised against INTS1 (A300-361A, Bethyl), INTS4 (A301-296A, Bethyl), INTS8 (HPA057299, Atlas Antibodies), INTS11 (A301-274A, Bethyl), GFP (sc-8334, Santa Cruz), PCNA (P8825, Sigma), RPB1 (14958, Cell Signalling Technologies), RPB1 Ser2P (3E10, Chromotek), RPB1 Ser5P (13523S, Cell Signalling Technologies), VCP (MA3-004, Thermo Scientific), and secondary HRP-conjugated antibodies were used. For immunocytochemistry, antibodies raised against PAX6 (901301, BioLegend), SOX2 (ab5603, Abcam), FOXG1 (ab18259, Abcam), TUJ1 (801201, BioLegend), MKI67 (550609, BD Pharmingen), MAP2 (188004, Synpatic Systems), SMI312 (837904, BioLegend), NESTIN (MAB1259, Merck), and Alexa fluorophore conjugated secondaries (Invitrogen) were used.

### Transfections

INTS8^ΔEVL^ NPC lines were created through CRISPR-Cas9 induced homology directed repair using a single stranded DNA oligonucleotide. Guide RNA GCCAGCTGTAACACTTCTTC, targeting exon 28 of INTS8, was cloned into vector pX330 eSpCas9-T2A-GFP. Together with the ssODN *G*TA CCA ACA AAA CTT AAT TTT TGA CAC ACC TCT AAT AAC ATC ACT TTTCTG TTT TGG AAG ATC AAA GCC ATC GGC CAG ACA GAG TTG AAT GCT AGC AAT CCA GAA CAG CTG GCA GCG CAG AGA AGG AAA AAA AAG TTT CTC CAA GCA ATG GCA AAA CTT TAC TTT TAA GCA GTT AAA TTT TTT TAA CTT TTA TTT *T*T, containing phosphorothioate linkages (*) at both ends, the vector was delivered to cells through nucleofection using the Amaxa Nucleofector kit V (VCA-1003). FACS-sorted GFP^+^ cells were seeded at single cell density, colonies isolated and genotyped for INTS8 c.2917-c.2925del by PCR and Sanger sequencing. Targeted regions were checked for large deletions (Table 1) and putative exonic off-target regions predicted by CRISPOR^59^ were verified by PCR and Sanger sequencing.

### Immunoprecipitation

Confluently grown NPCs were scraped in ice-cold PBS, nuclear extracts were prepared^60^ and diluted two-fold to 100 mM NaCl with C-0 (20 mM Hepes-KOH (pH7.6], 0.2 mM EDTA, 1.5 mM MgCl_2_, and 20% glycerol). Complete, EDTA-free protease inhibitor cocktail (Roche) was added to all buffers. Immunoprecipitation of INTS1 was performed in no-stick microtubes (Alpha Laboratories) from 200 µL NPC nuclear extract as previously described^61^, using 2.5 µg of INT1 antibody (Bethyl, A300-361A) or 2.5 µg GFP antibody (Santa Cruz, sc8334), 25U Benzonase (Novagen), and 25 µl protein A dynabeads equilibrated in C-100* (20 mM Hepes-KOH (pH7.6], 100 mM KCl, 0.2 mM EDTA, 1.5 mM MgCl_2_, 0.02% NP-40, and 20% glycerol) and blocked in 0.2 mg/ml chicken egg albumin (Sigma), 0.1 mg/ml insulin (Sigma), and 1% fish skin gelatin (Sigma) in C-100*. Beads were washed 4 times with C-100* buffer at 4 °C and proteins were eluted from the beads by boiling in 2x Laemmli Sample Buffer for 5 min at 95 °C.

### Flow cytometry analysis

Twenty-four hours prior labelling and/or harvesting, cells were seeded at a density of 45.000 cells/cm^2^. For Propidium Iodide Nucleic Acid staining, cells were harvested using accutase (Sigma A6964), washed, resuspended in 1ml ice cold PBS and fixed for 20 minutes following dropwise addition of 100% ice-cold EtOH. Cells were washed in ice-cold 1% BSA in PBS and stained for 45 minutes in the dark in 500 µl staining solution containing 30 µg/ml propidium iodide (Sigma P3566), 100 µg/ml DNase-free RNaseA in PBS. To quantify nascent transcription, cells were incubated with 1 mM EU (Axxora JBS-CLK-N0002 10) for 1 hour. Cells were harvested with accutase, transferred to a 96-well U-bottom plate, washed with PBS and fixed in 4% paraformaldehyde for 10 min at room temperature (RT). Cells were washed in 1% BSA PBS and permeabilized in 1X Click-iT saponin-based permeabilization buffer for 10 min on ice. Incorporated EU was labelled for 30 minutes at RT with a Click-iT reaction cocktail containing 2mg L-Ascorbic acid/ml (Sigma A5960), 2 mM CuSO4 and Alexa Fluor 647 Azide. After three washes with 1% BSA in PBS, cells were resuspended in 1% BSA in PBS containing 50 nM DAPI. Flow cytometry analysis was performed on the BD LSR Fortessa (BS Biosciences) and FlowJo was used for data analysis.

### Neural differentiation

Adherent cortical neuronal networks were generated from NPCs as described previously^26^. Briefly, sterile coverslips were coated with poly-L-ornithine (Sigma P4957) for 4h at RT, dried and incubated with a 100 µl drop of 50 µg/ml laminin solution (Sigma L2020) for at least 30 minutes at 37⁰C/5% CO2. To seed NPCs, the laminin droplet was replaced with a 100 µL droplet containing 200.000 cells in neural differentiation medium, consisting of neurobasal medium (Gibco 21103-049) containing 1% N2 supplement (Gibco 17502048), 2% B27-vitA supplement (Gibco 12587010), 1% MEM-NEAA (Gibco 11140-035), 2µg/ml laminin, 1% penicillin/streptomycin, and freshly added 20ng/µl BDNF (Prospec CYT-207), 20ng/µL GDNF (Prospec CYT-305), 1µM dibutyryl cyclic AMP (Sigma D0627), and 200µM ascorbic acid (Sigma A5960). Following cell attachment for 1h, 900µL of neural differentiation medium was added to the wells and medium was refreshed every second day.

### Immunocytochemistry

Cells were fixed in 4% paraformaldehyde in PBS for 10 minutes at RT, washed and permeabilized in 0.1% Triton X-100 in PBS. Coverslips were incubated with primary antibodies at 4⁰C overnight and with Alexa dye-conjugated secondary antibodies plus DAPI (Sigma, D9542) for 1h at RT, both diluted in blocking buffer (10% FBS / 0.1% Triton X-100 in PBS). Aqua-Poly/Mounted samples (Polysciences, 18606-20) were imaged with a Leica SP5 confocal microscope, processed and analysed with FIJI (ImageJ). Cell segmentation to allow quantification of nuclear Ki67 signal was performed with Cellpose^62^.

### RNA isolation and RT-qPCR

RNA was extracted using TRI reagent (Sigma-Aldrich T9424) and cDNA was generated the High-Capacity RNA-to-cDNA™ kit (Applied Biosystems 4387406) according to manufacturer’s instructions. RT-qPCRs were performed with 2 technical replicates and normalized to *NADH*, *SNAPIN* or both. For spike-in qPCR, 5% *Drosophila melanogaster* cells were added to a defined cell number in TRI reagent prior to RNA isolation.

### RNA sequencing

RNA was isolated from NPCs seeded at equal cell density using TRI reagent and purified on RNeasy columns (Qiagen). RNA quality was assessed on the Agilent bioanalyzer, all samples had RIN above 9.7. Sequencing libraries were prepared from 200 ng total RNA with the Illumina TruSeq Stranded mRNA Library Prep Kit and sequenced according to the Illumina TruSeq Rapid v2 protocol on an Illumina HiSeq2500 Sequencer for single 50bp reads, at least 21.5M reads per library. Adapter sequences were trimmed using in-house developed software and trimmed reads were mapped against the GRCh38 reference genome using HISAT2 (version 2.1.0^63^). Gene expression values were called using HTSeq-count (version 0.11.2^64^). Gene list was filtered to remove non-expressed genes after which differential expression analysis was performed using EdgeR. Differentially expressed genes were called with an FDR<0.05 and a |Log2FC ≥ 0.6|. Gene Ontology analysis was performed using Metascape^65^ and Gene Set Enrichment analysis was performed using GSEA software^29,66^.

### Chromatin immunoprecipitation

H9 NPCs suspended in PBS were crosslinked sequentially for 45 min with 2 mM disuccinimidyl glutarate (DSG, Thermo Fisher Scientific #20593) and for 10 min with 1% buffered formaldehyde solution (50 mM HEPES-KOH [pH 7.6], 100 mM NaCl, 1 mM EDTA, 0.5 mM EGTA, 11% Formaldehyde). Reactions were quenched with 125 mM glycine, chromatin was prepared, and ChIP performed as described^67^ using 150 µg chromatin as input, 10 µg of INTS11 antibody (Bethyl, A301-274A) or 10 µg of IgG antibody (Diagenode, C15410206) as control. Sequencing libraries were prepared from 10 ng ChIP DNA using the Thruplex DNA-Seq library prep kit (Takara Bio) and 16 PCR cycles. Libraries were sequenced on an Illumina Nextseq2000 sequencer. Paired-end clusters were generated of 50 bases in length, at least 50M read-pairs per sample. Adapters were trimmed using in-house developed software and reads were aligned to the GRCh38 reference genome using HISAT2 (version 2.1.0^63^). The alignments matching the reference genome were extracted from the BAM file and merged to continuous DNA fragments (represented in BED format). Oversized (>500bp) and duplicated DNA fragments were removed. Peaks were called using MACS2 callpeak^68^ using the BEDPE input format. Genome coverage bedgraphs were generated from the de-duplicated DNA fragments using bedtools^69^. Noise was deducted with MACS2 bdgcmp version 2.2.7.1^68^, using signal from input chromatin as control. The peaks across all samples were merged and quantified per sample. For annotation of INTS11 peak distribution, ChIPseeker v1.34.1 was used^70^. Heatmaps were generated using the deepTools2 package^71^ (version 3.5.1.0.1). Mean scores of 100 bp bins 4 kb around the peak summit were displayed.

### CUT&RUN

CUT&RUN was performed as described^72,73^. Briefly, 5*10^5^ cells were washed with wash buffer (20mM HEPES-KOH [pH7.5], 150mM NaCl, 0.5mM spermidine) and bound to Concanavalin A-coated magnetic beads activated in binding buffer (20mM HEPES-KOH pH7.5, 10mM KCl, 1mM CaCl_2_, 1mM MnCl_2_). Cells were permeabilized and incubated with 0.5 µg primary antibody for RNAPol2-CTD (8WG16, Sigma 05-952-I), 0.1 µg Ser5P D9N5I (CST 13523S) and 1 µg Ser5P 4H8 (Abcam ab5408) by rotation at 4⁰C overnight in 100µl wash buffer containing 0.05% digitonin and 2 mM EDTA. All steps were performed at 4⁰C with ice-cold buffers and in the presence of 1x complete EDTA-free protease inhibitor cocktail (Roche). Cells on beads were washed twice in wash buffer containing 0.05% digitonin (DigWash buffer) and incubated for 1h at 4⁰C in 100 µl DigWash buffer with 1 µg secondary antibodies Rabbit-a-Mouse (Abcam46540) or Guineapig-a-Rabbit (Abin101961) to enhance binding to ProtA. Samples were washed and incubated in 150 µl DigWash buffer containing 700 ng/ml ProtA-MNase fusion protein (kind gift from the Henikoff lab) for 1h at 4⁰C. Following two washes in DigWash buffer, samples were resuspended in 100 µl DigWash buffer and chilled to 0⁰C. MNase reactions were activated by the addition of 2 mM CaCl_2_ for 30 min at 0⁰C, stopped with STOP buffer (170 mM NaCl, 20 mM EGTA, 0.05% Digitonin, 18.75 mg/ml glycogen blue, 25 mg/ml RNase) and incubated at 37⁰C for 30 min. Alternatively, for total RNApol2 CUT&RUN, beads were washed twice in Low Salt rinse buffer (20mM HEPES-KOH pH7.5, 0.5mM spermidine, 0.05% digitonin), resuspended in 200µL incubation buffer (3.5mM HEPES-KOH pH7.5, 10 mM CaCl2, 0.05% digitonin) chilled to 0⁰C, and incubated for 30 min. DNA was extracted with Phenol-Chloroform-Isolamylalcohol (Sigma P3803) using MaXtract phase-lock tubes (Qiagen 129046). Libraries were prepared as described for ChIP, using 20 ng input DNA and 13 PCR amplification cycles, and sequenced on an Illumina Nextseq2500. Paired-end clusters were generated of 50 bases in length, at least 20M read-pairs per sample. Reads were processed, aligned and analyzed as described for ChIP. Pausing indices (PI) were calculated using the read density of both total RNAPol2 CUT&RUN datasets. Read counting was performed using the BedCoverage tool (Galaxy Version 2.0.2) and PIs were calculated by dividing length-normalised reads at TSS (± 250 bp) by length-normalised reads in the gene body (+500 bp – TES).

## Data availability

Next-generation sequencing data generated in this study is available from GEO.

### Author contributions

HBS: Methodology, Investigation, Formal analysis, Writing – review & editing. KNE, ALK, MPA, RMF, KK, TZ, ZA: Investigation. MRD, MF: Methodology. MH, WIJ: Formal analysis, Data curation, Software. DH, RAP: Writing – review & editing. DLCB: Conceptualization, Methodology, Writing – original draft, Supervision, Funding acquisition.

## Acknowledgements

DLCB is supported by a Horizon 2020 Marie Curie Individual Fellowship (#799214) and an Erasmus University Rotterdam Fellowship.

