## Supplemental_figures_methods for "The Integrator complex prevents premature neuronal differentiation through global control of transcription elongation"

**A**

| Off-target gene | Genomic location off-target site | CFD score | FW primer | RV primer |
|---|---|---|---|---|
| *PCOLCE2* | chr3:142,535,376-142,535,398 | 0.21 | GACAAACCAGTGCCACACAC | AACGACAAAATTGGGCTGAC |
| *ZNF581* | chr19:56,156,183-56,156,205 | 0.21 | GGTGTCCCCTACACAGTGCT | CCAGGTGTCTGGACAGGAGT |
| *PCYOX1L* | chr5:148,744,278-148,744,300 | 0.19 | AGTCGCTTAACAAGGGCTGA | AGGGAGGGAACAGACCACTT |
| *RNU7-25P* | chr18:3,093,559-3,093,581 | 0.16 | CCACAAACCACACCTCCTCT | GTCCCAATGGCTCTGAAAAA |
| *JMJD1C* | chr10:64,936,056-64,936,078 | 0.16 | GGCAGACTGGTGCAGGTAAT | GTGGAAGGAGGTGGTGAGAA |
| *CACNA1G* | chr17:48,703,704-48,703,726 | 0.14 | GCTAGTCCATGCTGGAGAGG | AGCTGACGGCGATAGAGTGT |
| *CHML* | chr1:241,797,274-241,797,296 | 0.10 | AGGTCCAAAACAGCATTTCG | ACCATGACATGCATGAAGGA |
| *BCL2L15* | chr1:114,424,465-114,424,487 | 0.09 | AGGGAAGCCTGTTTTCCTGT | AGCCTCCTGCACTTTTTGAG |
| *ARHGEF7* | chr13:111,775,058-111,775,080 | 0.07 | TTTGCAGGATGGTGTCATGT | CCCTTCTCCACTCTGCTTTG |
| *GSN* | chr9:124,091,425-124,091,447 | 0.06 | ACGGCTGAAGGACAAGAAGA | GTACCGCCACTCCCAGTTTA |

**B**

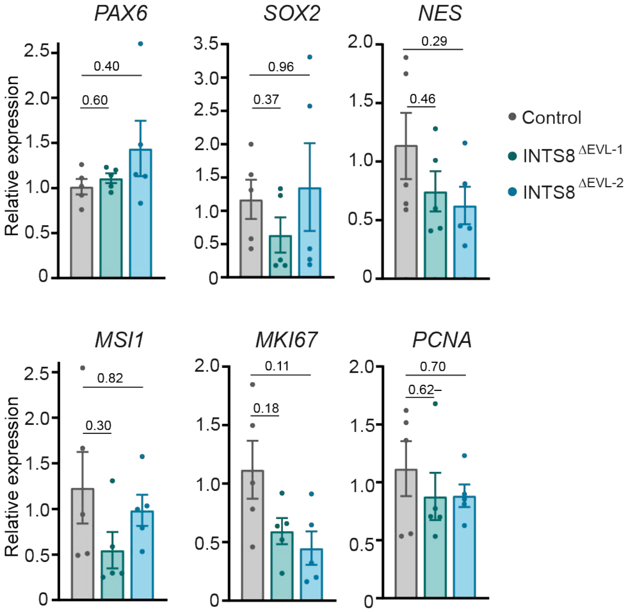


**C**

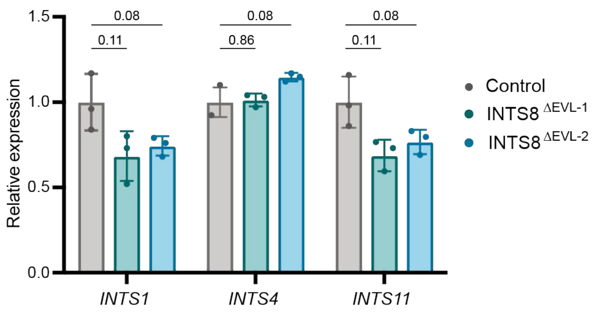


Figure S1 Somsen et al.

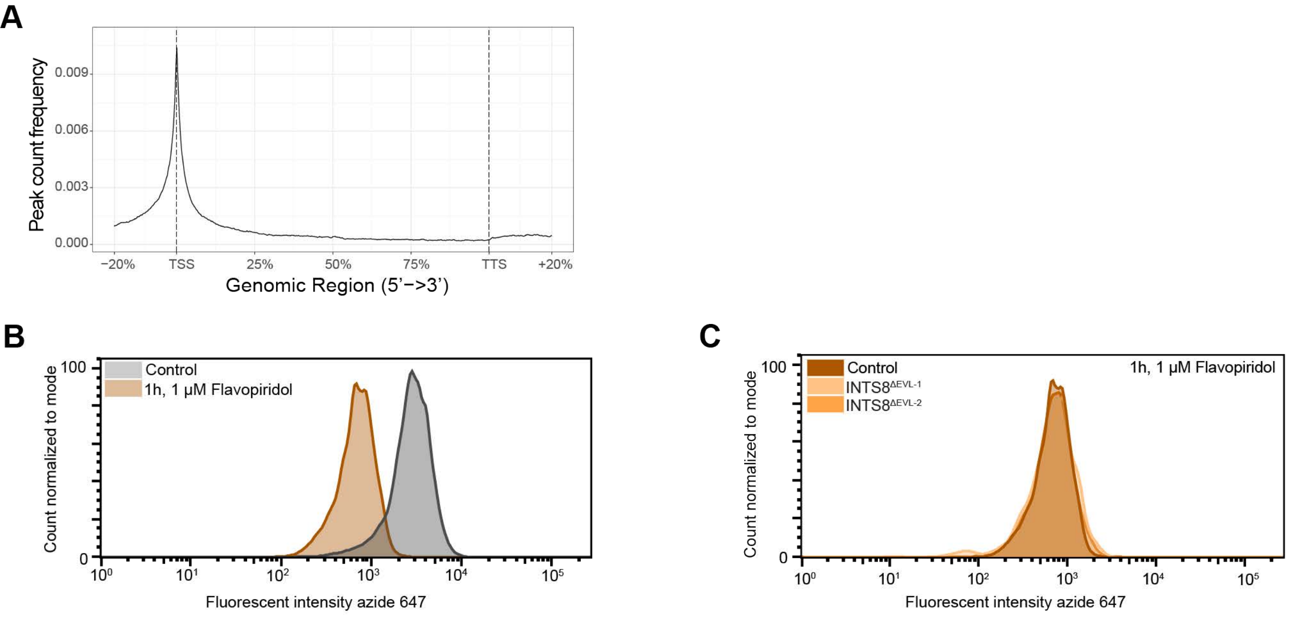


Figure S2, Somsen et al.

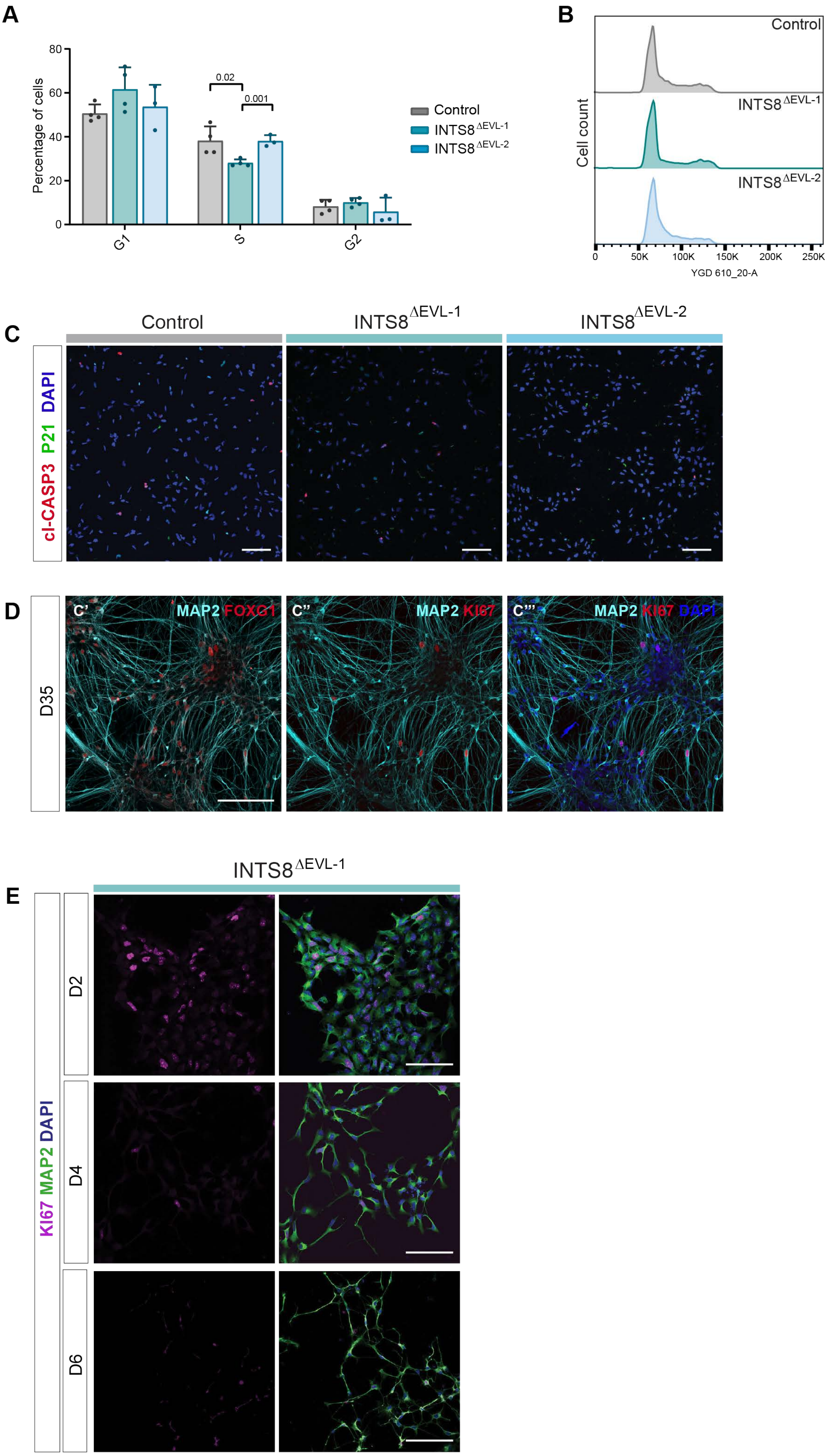


Figure S3, Somsen et al.

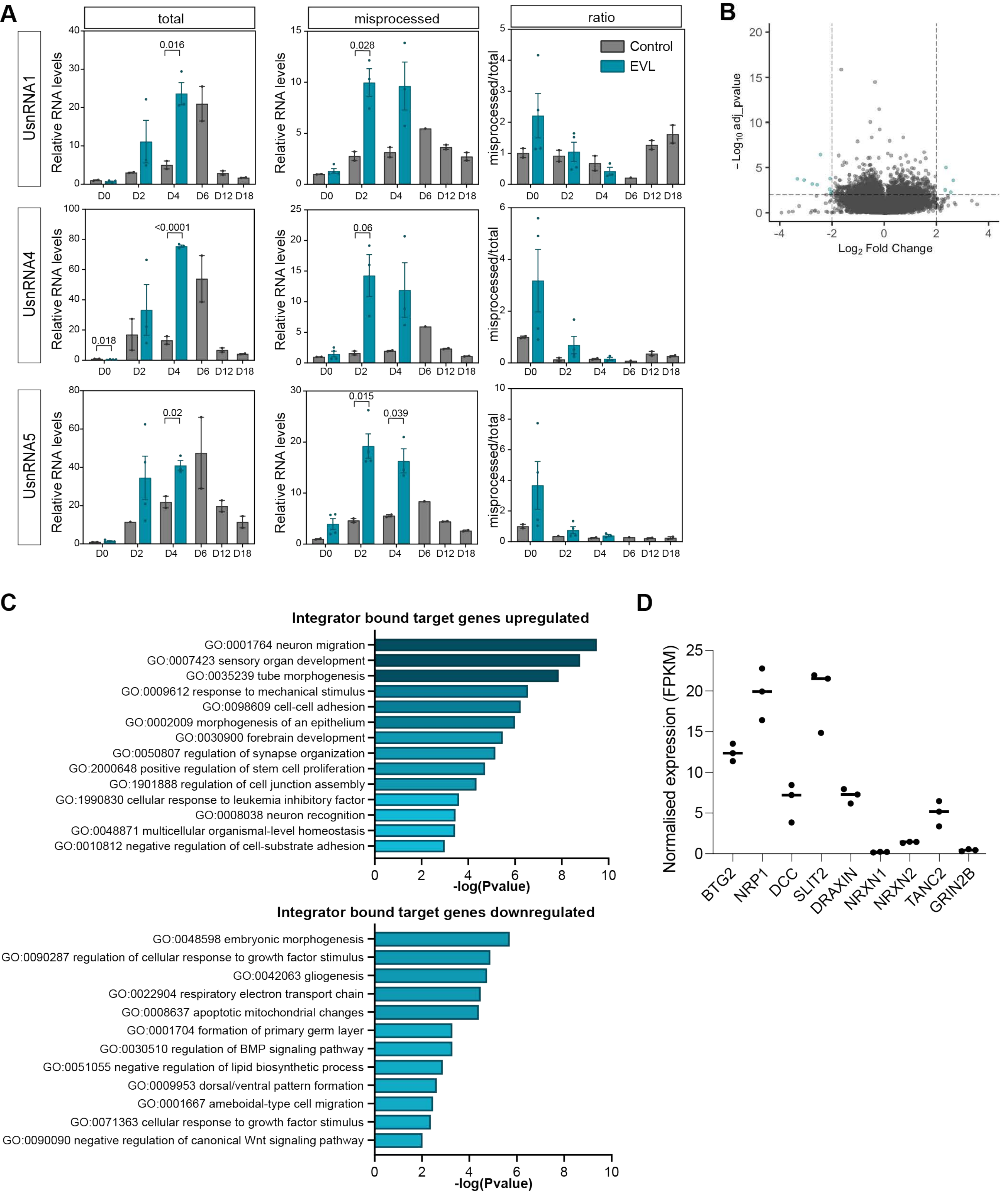


Figure S4, Somsen et al.

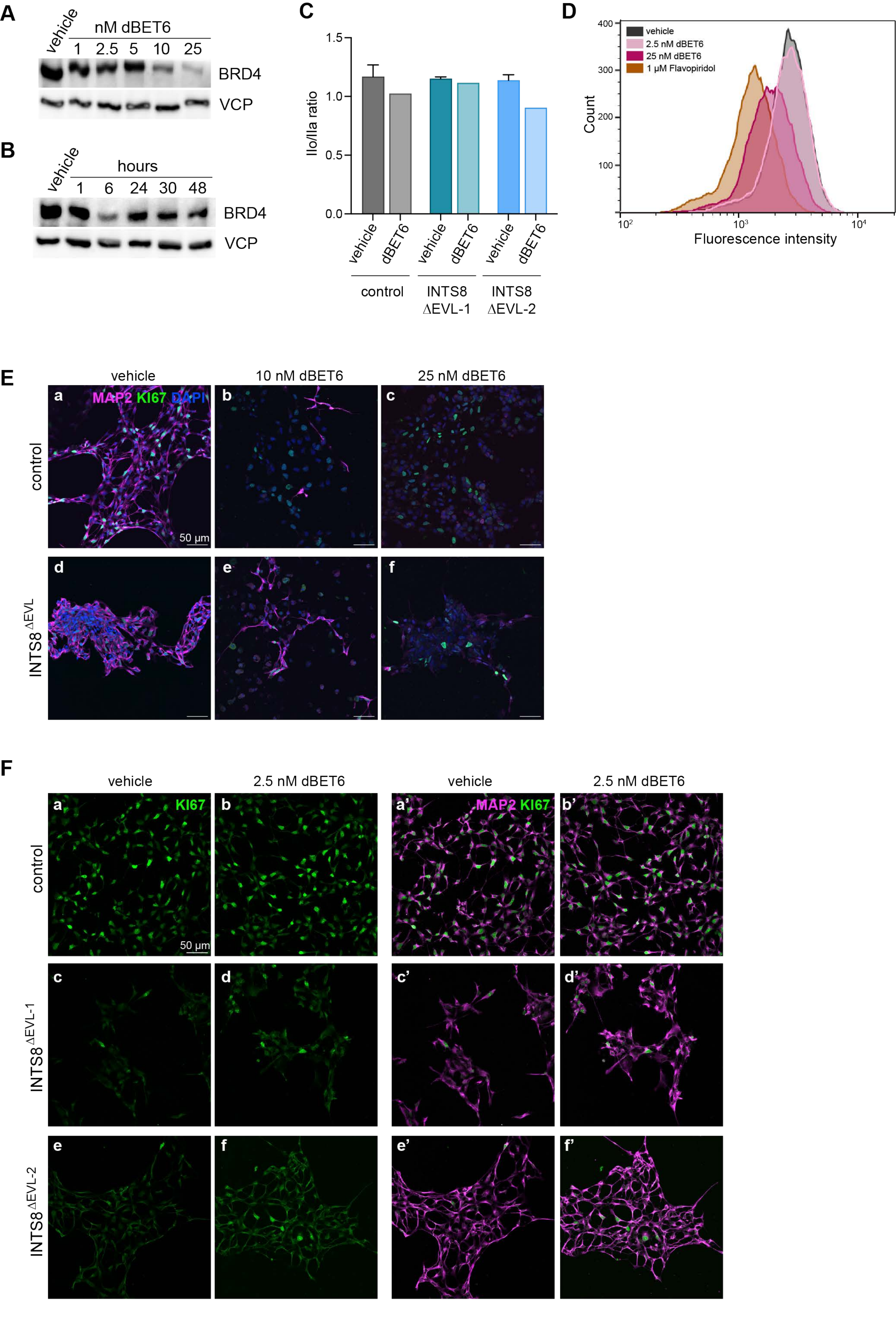


Figure S5, Somsen et al.

### SUPPLEMENTAL METHODS

#### Data analysis

To detect differences in exon usage we used DEXSeq (version 1.28.1[1,2]). Briefly, exon counting bins were defined based on a flattened gene model and used to count aligned reads per bin for each sample. We then performed a standard differential exon usage analysis and called differentially used exons based on |Log2FC ≥ 2| and $p_{adj} \leq 0.01$).


1 Anders, S., Reyes, A. & Huber, W. Detecting differential usage of exons from RNA-seq data. *Genome Res* **22**, 2008-2017 (2012). https://doi.org/10.1101/gr.133744.111

2 Reyes, A. *et al.* Drift and conservation of differential exon usage across tissues in primate species. *Proc Natl Acad Sci U S A* **110**, 15377-15382 (2013). https://doi.org/10.1073/pnas.1307202110


**Table 1. Quantitative PCR primers**

| Target | FW primer | RV primer |
|---|---|---|
| *PAX6* | TTTGCCCGAGAAAGACTAGC | CATTTGGCCCTTCGATTAGA |
| *SOX2* | GGGAAATGGGAGGGGTGCAAAAGAGG | TTGCGTGAGTGTGGATGGGATTGGTG |
| *NES* | CAGCGTTGGAACAGAGGTTGG | TGGCACAGGTGTCTCAAGGGTAG |
| *MSI1* | TGACCAAGAGATCCAGGGGT | CGATTGCGCCAGCACTTTAT |
| *MKI67* | GATCGTCCCAGTGGAAGAG | AAGGCCAGGTATAATCCGTA |
| *PCNA* | CTCTTCCCTTACGCAAGTCT | GAAAGTCTAGCTGGTTTCGG |
| *UsnRNA1* total | ATACCATGATCACGAAGGTGGTT | CAGTCCCCCACTACCACAAATTA |
| *UsnRNA1* misprocessed | TACCTGGCAGGGGAGATACC | TACCTGGCAGGGGAGATACC |
| *UsnRNA4* total | GCAGTATCGTAGCCAATGAGGTCTA | CCAGTGCCGACTATATTGCAAGTC |
| *UsnRNA4* misprocessed | CGTAGCCAATGAGGTCTATCCG | CCTCTGTTGTTCAACTGCAAGAAA |
| *UsnRNA5* total | CTCTGGTTTCTCTTCAGATCGCA | TTGGGTTAAGACTCAGAGTTGTTCC |
| *UsnRNA5* misprocessed | ATTTCCGTGGAGAGGAACAACTC | GCACCATTGAACAGAAAAAGGAA |
| *BTG2* | CATCATCAGCAGGGTGGC | CCCAATGCGGTAGGACAC |
| *NRP1* | GTCTTCAGGGCCATTTCTTT | GTTCTGGGAACATTCAGGAC |
| *DCC* | TCCAACAATCCTGCTGTCGT | CTTCTTCCTGCTCCGAAACCT |
| *SLIT2* | GAATTTGTCTGCAGTGGTCA | TGGAAGATTTGTGGGGATCT |
| *DRAXIN* | CGACTGGACCGATTATGAAGAC | CGGCTGGTGATGTTTCGTTAC |
| *NRXN1* | ACAACAATGTGGAAGGTCTG | TGCTTTGAATGGGGTTTTGA |
| *NRXN2* | GTGTCCAAGCGATGATGAG | GCAATATTAACTCTCCTCCAGT |
| *TANC2* | AAGAAGTTAACTGCCCCTCT | GTTTCTGAGCTGAAGTGAGG |
| *GRIN2B* | GGCTTCTTCACCTCTAGGAA | AAAGGATATGCATTCGGACG |
| *NADH* | ACTGGCTACTGCGTACATCC | AGATGCGCCTATCTCTTTCC |
| *SNAPIN* | AGCTCGACTCTCACGTACAC | GCCGGGCATTAAGTAGCTTC |
| *Dm_Act42A* | CTAAGCAGTAGTCGGGCTGG | GTCTGCAATGGGTGTGTTCG |

### SUPPLEMENTAL FIGURE LEGENDS

**Figure S1, related to Figure 1. Characterisation of INTS8$^{\Delta EVL}$ NPC lines.**

(A) CFD scores as determined by CRISPOR for top 10 predicted exonic off-target locations for CRISPR-Cas9 hybrid. Primers used for genotyping by Sanger sequencing are indicated. (B) Mean relative expression values of indicated genes as determined by RT-qPCR analysis. Error bars represent SEM, n=5, p-values determined by Brown-Forsythe and Welch ANOVA followed by Dunnett's T3 multiple comparisons test. (C) Mean relative expression values of indicated genes as determined by RT-qPCR analysis. Error bars represent SEM, q-values determined by two-stage step-up method of Benjamini, Krieger and Yekutieli, n=3.

**Figure S2, related to Figure 2. Integrator as global attenuator of transcription elongation**. (A) Profile of peak distribution over scaled gene body regions. TSS, transcription start site; TTS, transcription termination site. (B) Histogram from flow cytometry showing incorporated EU fluorescence intensity in vehicle-treated NPCs or NPCs treated for 1 hour with 1 µM Flavopiridol to inhibit transcription elongation. (C) Histogram from flow cytometry showing incorporated EU fluorescence intensity in different NPC lines treated for 1 hour with 1 µM Flavopiridol to inhibit transcription elongation.

**Figure S3, related to Figure 3. Cell cycle parameters or cell viability are not affected in INTS8$^{ΔEVL}$ NPCs.** (A) Distribution of cells over the different cell cycle phases as determined by propidium iodide-labelling and flow cytometry analysis. Error bars represent SEM, n=4. P-values determined with unpaired Student's t test. (B) Representative histograms showing PI fluorescence intensity (x-axis) in indicated NPC lines. (C) ICC for cleaved caspase-3 and p21 (CDKN1A) in indicated NPC lines. Scale bar represents 100 µm. (D) ICC for MAP2, KI67 and FOXG1 on control NPCs at D35 of neural differentiation. Scale bar represents 100 µm. (E) Differentiating NPCs subjected to ICC with KI67 and MAP2 antibodies at indicated timepoints during neuronal differentiation. Scale bar xxx µm.

**Figure S4, related to Figure 4 and 5. Impact of INTS8$^{ΔEVL}$ variant on UsnRNA processing, differential exon and gene expression.**

(A) RT-qPCR analysis of total and misprocessed UsnRNA transcripts at different timepoints of neural differentiation. Error bars represent SEM, n=2 for control, n=4 for INTS8$^{ΔEVL}$. P-values determined with unpaired Student's t test. (B) Volcano plot visualizing differentially expressed exons (green dots) as determined by RNA-seq (|$Log_2$FC>2|, FDR <0.01). (C) Metascape GO analysis of DEGs containing bound by INTS11 in their promoter region. (D) Normalised gene expression (FPKM) for DEGs in control NPCs.

**Figure S5, related to Figure 6. Data supporting use of EU and dBET6 to monitor and modulate nascent transcription.**

(A) Western blot for BRD4 on NPCs treated for 1 hour with vehicle or indicated doses dBET6. VCP was used as loading control. (B) Western blot for BRD4 on NPCs treated with 2.5 nM dBET6 for indicated duration. VCP was used as loading control. (C) Quantification of ratio between hyper- (IIo) and hypophosphorylated (IIa) RPB1 in NPCs treated for 24 hrs with vehicle or 25 nM dBET6. (D) Flow cytometry histogram showing effect of indicated dBET6 doses on nascent transcription as measured by EU incorporation. (E) Immunocytochemistry with indicated antibodies on NPCs at neural differentiation D4, treated with vehicle (a, d), 10 nM (b, e), or 25 nM (c, f) dBET6 for 96 hours. Scale bar represents 50 µm. (F) Additional images of indicated NPC lines at D4 of in vitro differentiation treated with vehicle or 2.5 nM dBET6 for 48-96 hours. Scale bar represents 50 µm.
